# The international sinonasal microbiome study (ISMS): a multi-centre, multi-national collaboration characterising the microbial ecology of the sinonasal cavity

**DOI:** 10.1101/548743

**Authors:** Sathish Paramasivan, Ahmed Bassiouni, Arron Shiffer, Matthew R Dillon, Emily K Cope, Clare Cooksley, Mahnaz Ramezanpour, Sophia Moraitis, Mohammad Javed Ali, Benjamin Bleier, Claudio Callejas, Marjolein E Cornet, Richard G Douglas, Daniel Dutra, Christos Georgalas, Richard J Harvey, Peter H Hwang, Amber U Luong, Rodney J Schlosser, Pongsakorn Tantilipikorn, Marc A Tewfik, Sarah Vreugde, Peter-John Wormald, J Gregory Caporaso, Alkis J Psaltis

**Affiliations:** Department of Otolaryngology, Head and Neck Surgery, University of Adelaide, Adelaide, Australia; Pathogen and Microbiome Institute, Northern Arizona University, Arizona, USA; Dacryology Service, LV Prasad Institute, Hyderabad, India; Department of Otolaryngology, Massachusetts Eye and Ear Infirmary, Harvard Medical School, Boston, USA; Department of Otolaryngology, Pontificia Universidad Catolica de Chile, Santiago, Chile; Department of Otorhinolaryngoloy, Amsterdam UMC, Amsterdam, The Netherlands; Department of Surgery, University of Auckland, Auckland, New Zealand; Department of Otorhinolaryngology, University of Sao Paulo, Sao Paulo, Brazil; Department of Otolaryngology, Rhinology and Skull base, University of New South Wales, Sydney, Australia; Faculty of Medicine and Health sciences, Macquarie University, Sydney, Australia; Department of Otolaryngology -Head and Neck Surgery, Stanford University, Stanford, California, USA; Department of Otolaryngology -Head and Neck Surgery, University of Texas, Texas, USA; Department of Otolaryngology, Medical University of South Carolina, Charleston, South Carolina, USA; Department of Otorhinolaryngology, Faculty of Medicine, Siriraj Hospital, Mahidol University, Bangkok, Thailand; Department of Otolaryngology - Head and Neck Surgery, McGill University, Montreal, Canada

**Keywords:** Chronic rhinosinusitis, microbiota, next-generation sequencing, 16S rRNA gene, sinus, microbiome, polyp

## Abstract

The sinonasal microbiome remains poorly defined, with our current knowledge based on a few cohort studies whose findings are inconsistent. Furthermore, the variability of the sinus microbiome across geographical divides remains unexplored. We characterise the sinonasal microbiome and its geographical variations in both health and disease using 16S rRNA gene sequencing of 410 individuals from across the world. Although the sinus microbial ecology is highly variable between individuals, we identify a core microbiome comprised of *Corynebacterium, Staphylococcus, Streptococcus, Haemophilus*, and *Moraxella* species in both healthy and chronic rhinosinusitis (CRS) cohorts. *Corynebacterium* (mean relative abundance = 44.02%) and *Staphylococcus* (mean relative abundance = 27.34%) appear particularly dominant in the majority of patients sampled. There was a significant variation in microbial diversity between countries (p = 0.001). Amongst patients suffering from CRS with nasal polyps, a significant depletion of *Corynebacterium* (40.29% vs 50.43%; p = 0.02) and over-representation of *Streptococcus* (7.21% vs 2.73%; p = 0.032) was identified. The delineation of the sinonasal microbiome and standardised methodology described within our study will enable further characterisation and translational application of the sinus microbiota.

## MAIN TEXT

The important role of human microbiota in both health and disease has become increasingly recognised. Microbial communities encode millions of genes and associated functions which act in concert with those of human cells to maintain homeostasis.^1^ Numerous studies have now established the microbiota as an important contributor to essential mammalian functions such as metabolism^2^, biosynthesis^3^, neurotransmission^4,5^ and immunomodulation^6,7^. Characterizing the composition and diversity of normal, healthy microbial communities is a cornerstone to developing our understanding of dysbiosis, pathophysiology and, ultimately, directing therapy. To this end, advent of next-generation sequencing (NGS) has revolutionised our appreciation of the host microbiota and its polymicrobial nature.^8,9^

In many cases the entire microbial community – commensal, symbiotic, pathogenic bacteria, fungi, archaea, and viruses – play critical roles in both health and disease pathogenesis. The host-microbiota interface is particularly important in chronic mucosal inflammatory conditions where the microbiota interact directly with the host. These conditions are often poorly understood, multifactorial in nature, have heterogeneous clinical presentations and vary in treatment response.^10,11^ Furthermore, single causative pathogens are rarely identified, and culture directed-antibiotics often fail to demonstrate efficacy.^12^ It is plausible that a better understanding of the microbiome of such conditions may be key to unraveling their underlying pathogenesis.

The sinonasal mucosa is continuously exposed to external particulate matter and microbes, but it is relatively immunodeplete with no native secondary lymphoid organ systems.^13^ The sinus microbiota is thought to play key roles in multiple extra-nasal conditions, such as providing a nidus of recurrent infection in cystic fibrosis patients^14^, and otitis media.^15^ In addition, there is evidence of microbial influences in the development, progression and severity of chronic rhinosinusitis (CRS).^16^ This multifactorial condition, with an estimated world-wide prevalence of approximately 10%,^11^ represents one of the most common diagnoses for inappropriate antibiotic prescription and is a source of significant morbidity and healthcare costs.^11,17–19^ To date, despite a number of well-designed research efforts to define the nature of the sinonasal microbiome and its role in CRS pathogenesis, many uncertainties persist. This is in part due to the difficulty in bacterial collection from the nose itself. Unlike the gut and oral cavity where the bacterial burden is high and access to microbiota relatively easy (either via faecal samples or oral wash),^20,21^ the sinonasal tract has a low microbial burden and access is difficult due to both the narrow nasal orifice and discomfort for the awake patient. Therefore, a replicable sample with appropriate bacterial abundance for 16S rRNA gene sequencing is currently attainable only during nasal surgery. To date, the majority of published studies have been small in size, with heterogeneous patient populations and inconsistency in collection methods, sample site, processing techniques and bioinformatics pipelines.^22^ Ultimately, the consequence has been non-comparable results with no universal consensus on the constituents of the healthy sinonasal microbiota or the dysbiosis that occurs in disease.^23–29^

To address these limitations, we investigate the sinonasal microbiome using 16S rRNA gene-sequencing on a large, multi-centre, international cohort implementing consistent sampling, processing and bioinformatics methods. We aim to (1) characterise the normal sinonasal microbiome, (2) assess for any geographical or clinical influences and (3) identify any changes associated with CRS within and across geographical sites.

## RESULTS

### Patient cohort

Middle meatus specimens were collected for 16S rRNA gene sequencing (V3-V4 hypervariable region) on the Illumina MiSeq platform (see Methods). Thirteen centres, across five continents, participated in patient sampling. 532 samples (194 healthy controls and 338 CRS patients) successfully went through all stages of transport and processing to be sent for sequencing. High-quality sequences were analysed using QIIME 2.^30^ A total of 410 patients, aged between 20 and 75, reached the final stage of analysis. This population included 139 non-CRS healthy controls, 99 patients without nasal polyposis (CRSsNP) and 172 CRS patients with nasal polyposis (CRSwNP). Supplementary Figure S1 demonstrates sample distribution by centre.

### The sinonasal microbiome in healthy sinuses is dominated by Corynebacterium and Staphylococcus

We first investigated the composition of the healthy sinonasal microbiome by intra-operatively sampling the 139 non-CRS control patients. (See Methods) Our analysis demonstrated the dominance of *Corynebacterium* (mean relative abundance = 48.7%; prevalence = 88.49%) and *Staphylococcus* species (mean relative abundance = 29.25%; prevalence = 79.86%) in the sinonasal microbiome of healthy patients. These were both the most abundant and prevalent genera amongst our population (Table 1). This finding has been observed in some but not all previously reported studies with variability in sampling and analysis techniques likely accounting for such discrepancies.^22^

**Table 1:** Abundance and prevalence of genera found in microbiota of healthy non-CRS patients.

| Genera | Mean Relative Abundance (%) | Prevalence (%) |
| --- | --- | --- |
| *Corynebacterium* | 48.7 | 88.49 |
| *Staphylococcus* | 29.25 | 79.86 |
| *Moraxella* | 3.86 | 12.23 |
| Streptococcus | 2.81 | 20.86 |
| Haemophilus | 2.23 | 12.95 |
| unidentified<br>(Enterobacteriaceae) | 2.16 | 13.67 |
| Serratia | 1.79 | 2.16 |
| Alloiococcus | 1.61 | 20.86 |
| unidentified (Neisseriaceae) | 1.25 | 12.23 |
| Pseudomonas | 0.75 | 2.88 |

### Microbiome composition is altered in CRSwNP and by geographical location

To explore influences of microbial composition, we examined the taxonomic profiles of our patient cohort once grouped by (1) disease cluster and (2) centre of origin (Figure 1) utilizing mixed modeling to control for the “centre” variable by assigning it as a random effect (See Methods). CRSsNP patients demonstrated no significant differences in the relative abundance of the top ten most abundant organisms when compared to healthy controls. (p > 0.05; mixed model analysis) The relative abundance of most organisms also remained stable between controls and CRSwNP but for two genera (Figure 1A): *Corynebacterium* was found to be significantly reduced in CRSwNP when compared to controls (40.29% vs 50.43%; mixed model analysis; p = 0.02) whilst *Streptococcus* was increased (7.21% vs 2.73%; mixed model analysis; p = 0.032). Interestingly, the most commonly cultured pathogens in CRS, *Staphylococcus* and *Pseudomonas* were similar between all cohorts.

**Figure 1:**
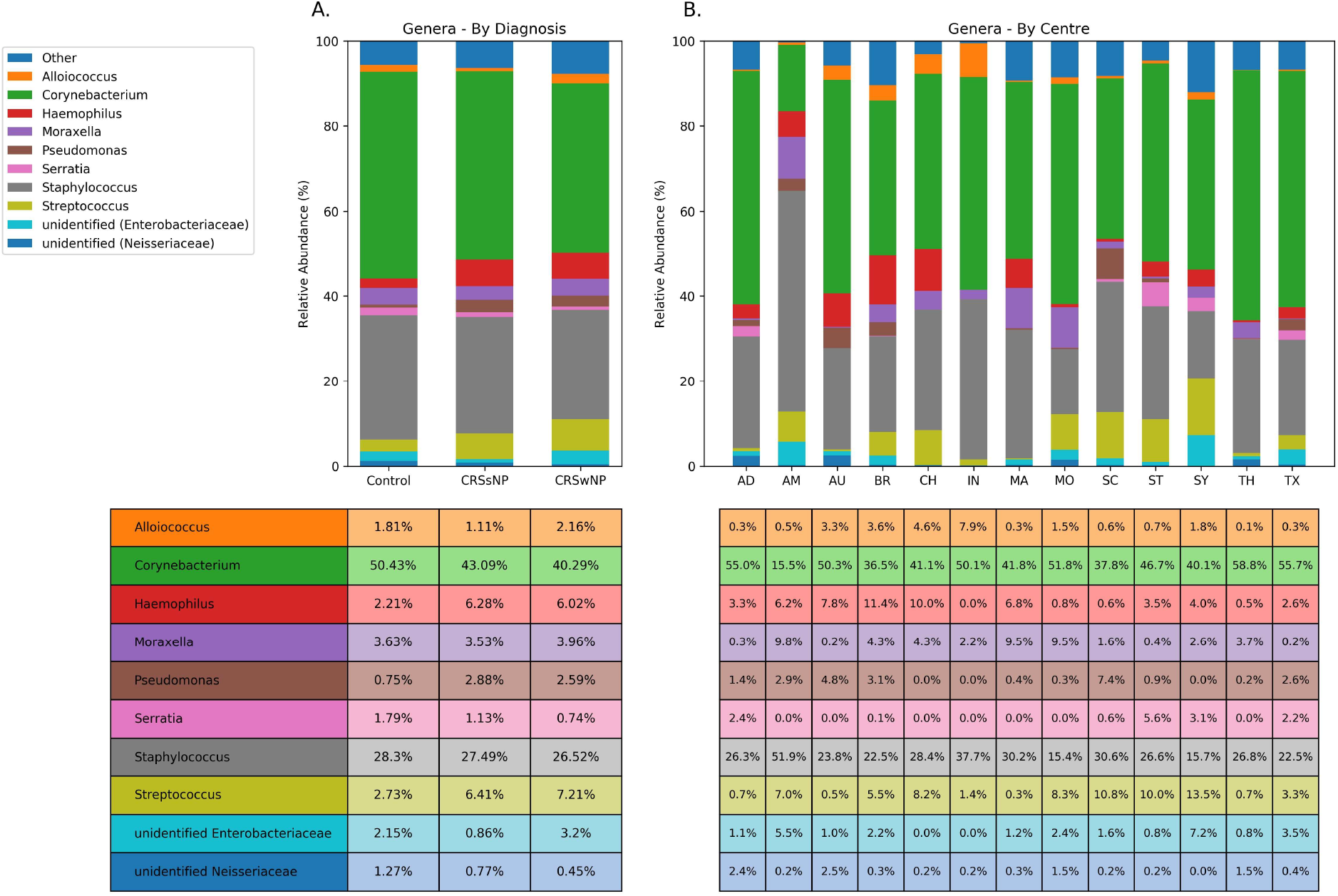
Microbiome taxonomic profiles by disease status and centres. Sinonasal microbial composition of patients (n = 409) when grouped by disease and centres. Accompanying tables demonstrate the corresponding relative abundances of the top ten most abundant organisms found within our cohort. The abundances of genera in A (by disease status) have been adjusted according to the mixed model that accounted for the centre as a random variable. CRSsNP = Chronic rhinosinusitis without nasal polyps; CRSwNP = chronic rhinosinusitis with nasal polyps; Control = Healthy, non-CRS patients AD = Adelaide; AM = Amsterdam; AU = Auckland; BR = Brazil; CH = Chile; IN = India; MA = Massachusetts; MO = Montreal; SC = South Carolina; ST = Stanford; SY = Sydney; TH = Thailand; TX = Texas.

By contrast, comparisons between centres revealed a higher degree of variability in the microbiome composition (Figure 1B). Samples from Amsterdam were significantly different from the remainder of the cohort, with increased *Staphylococcus* (51.94%) and marked depletion of *Corynebacterium* (15.51%).

Amongst the remaining centres, each appeared to have some variability, with individual regions displaying increased or decreased relative abundance in specific taxa. *Streptococcus*, for example, made up 13.48% of the Sydney microbiome, but was almost absent amongst the Adelaide (0.65%), Auckland (0.45%), Massachusetts (0.32%) and Thailand (0.72%) cohorts. Similar variability can be seen for almost all bacterial taxa examined (Figure 1B). While centres’ samples were appeared fairly similar in microbial composition, these results would suggest that centre-specific microbiome profiles can be found. This variability may account for some of the inconsistencies observed in the literature, particularly between findings from different institutes.

### Microbial diversity is influenced by geographical location

Alpha diversity amongst cohorts was performed utilising Faith’s Phylogenetic Diversity (PD) index.^31^ Comparison between disease states demonstrated no significant differences between controls, CRSsNP and CRSwNP (Figure 2A). Overall, alpha diversity was significantly different between centres (Kruskall-Wallis; p < 0.001; Figure 2B). Of particular interest was the finding of a significantly lower alpha diversity for samples from Amsterdam compared to the other centres (mean PD = 1.27, p < 0.01). This may be related to the compositional findings of high staphylococcal abundance in Amsterdam.

**Figure 2:**
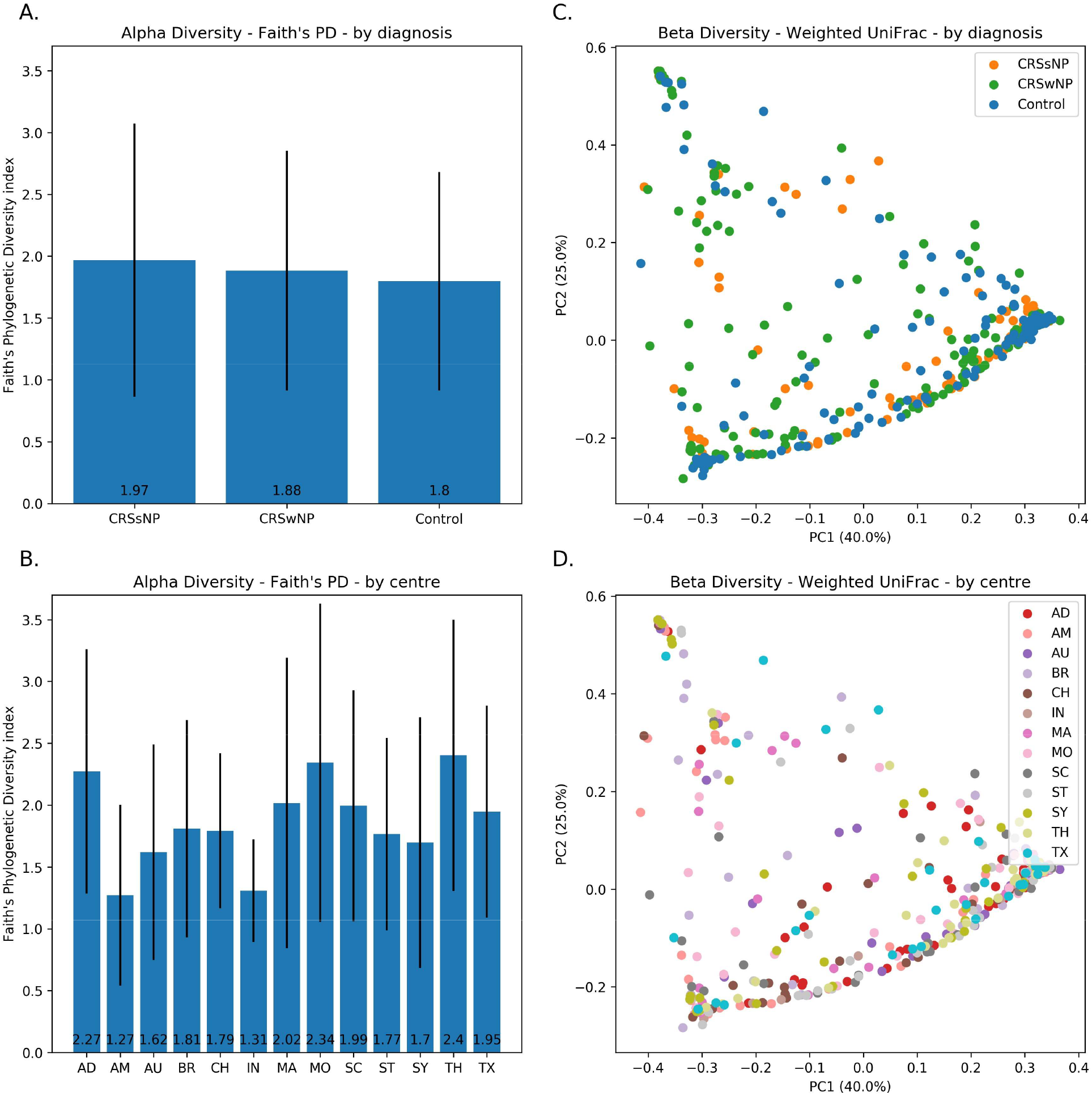
Alpha and beta diversity plots. Alpha diversity, derived from Faith’s Phylogenetic Diveristy Index, demonstrated for this cohort (n = 409) when grouped by disease and by collection centre. Error bars represent 95% confidence intervals. Beta diversity is demonstrated here as a Principal Coordinate Analysis (PCoA) plot of the Weighted-UniFrac distance matrix. Each dot represents a single patient. Similarities between patients are represented by their proximity to each other on the graph. Again, patients are classified by disease and centre. Component 1 (PC1) is represented on the x-axis and component 2 (PC2) on the y-axis. Patients tended towards clustering into one of three groups, as visualised. PD = Phylogenetic diversity CRSsNP = chronic rhinosinusitis without nasal polyps; CRSwN p= chronic rhinosinusitis with nasal polyps; Control = healthy, non-CRS patients AD = Adelaide; AM = Amsterdam; AU = Auckland; BR = Brazil; CH = Chile; IN = India; MA = Massachusetts; MO = Montreal; SC = South Carolina; ST = Stanford; SY = Sydney; TH = Thailand; TX = Texas.

Multivariate analysis on the beta diversity distance matrix was done using PERMANOVA (with 999 permutations) to explore the significance of the diagnosis versus the centre variables in a single model. This showed a significant effect of the centre covariate (pseudo-F = 2.51; p = 0.001) on the weighted UniFrac^32^ distances, while the diagnosis covariate was not significant (pseudo-F = 1.66; p = 0.1).

We also performed Principal Coordinates Analysis (PCoA) on the Weighted-UniFrac distance matrix. This did not show clustering by disease state (Figure 2C and 2D), although the plot may suggest a distribution amenable to unsupervised clustering. Patients tended to cluster into three groups on a continuum on the PCoA. These individual clusters could represent specific microbial community types, similar to what has been previously reported by Cope et al.^23^ Investigation into the constitution of these groupings, and their association with underlying clinical or pathological factors in a large multi-institutional cohort remains a topic of future investigation.

### The core sinonasal microbiome is composed of five genera

We defined a core sinonasal microbiome by analysing the most abundant organisms along with their cumulative prevalence. This was performed across all samples and also across the three different disease groups. The results of this investigation confirmed a high prevalence of the top five abundant genera (*Corynebacterium, Staphylococcus, Streptococcus, Haemophilus*, and *Moraxella*), which together reached a cumulative prevalence in 98-100% of samples in all patient groups. Presence of at least one of these taxa in nearly all patients suggests that they make up the core sinonasal microbiome, regardless of disease status (Figure 3). Amongst control patients, *Corynebacterium* and *Staphylococcus* were present in 98.6% of patients, again suggesting a likely key commensal function of these two genera in the healthy state.

**Figure 3:**
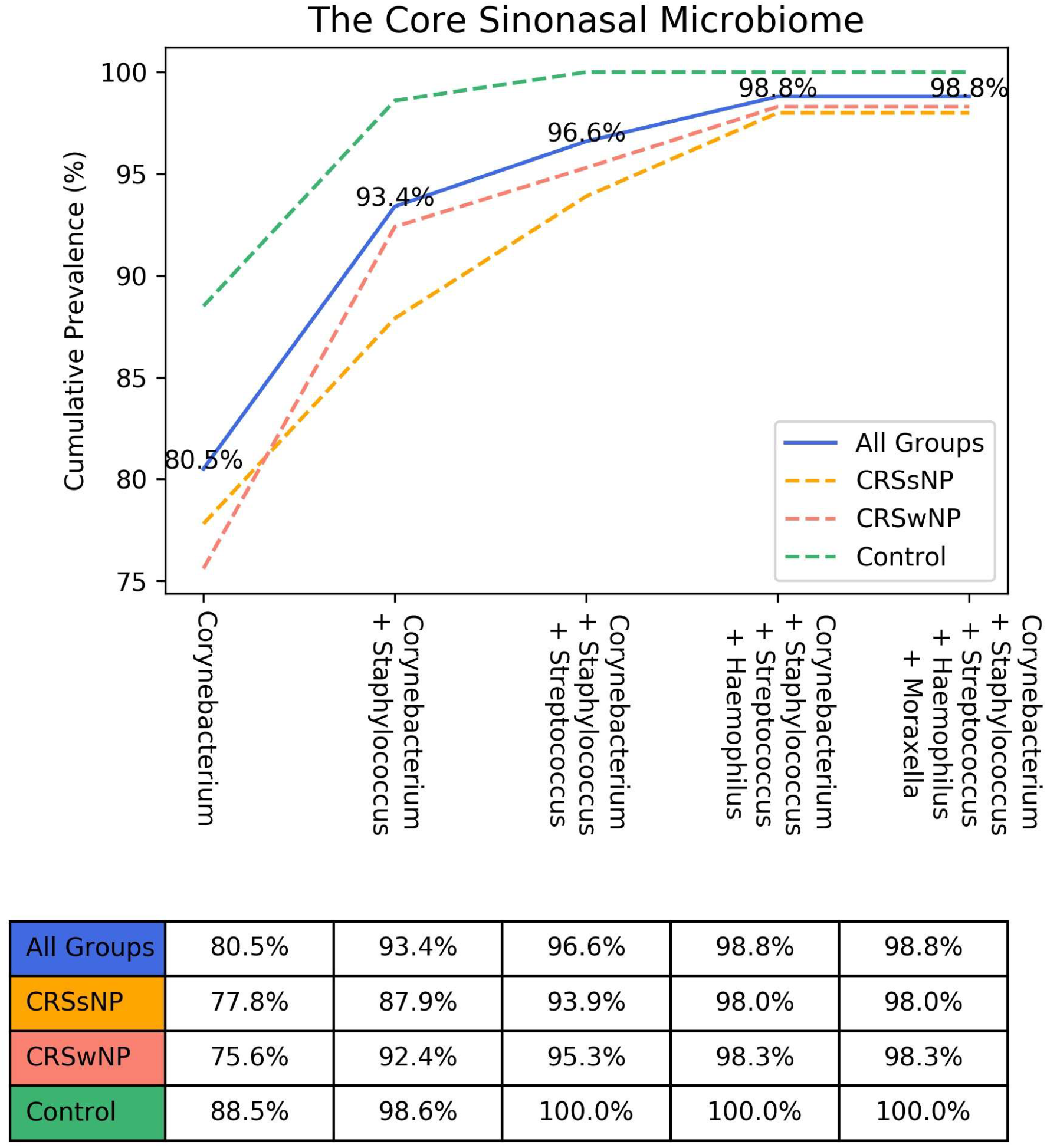
Cumulative microbial prevalence for the top abundant organisms. Cumulative prevalence for the top five most abundant organisms is presented above. For all groups, we commence with the most abundant organism then add subsequent microbial taxa in a descending fashion based on relative abundance. CRSsNP = chronic rhinosinusitis without nasal polyps; CRSwNP = chronic rhinosinusitis with nasal polyps; Control = healthy, non-CRS patients.

### Clinical covariates do not correlate with changes to taxonomic composition

To determine whether any host factors may contribute to the stability of the sinonasal microbiome, patients were separated by known clinical variables that contribute to CRS (Figure 4). There were no significant differential abundances (of the 10 topmost abundant genera) for all six clinical covariates examined (asthma, aspirin sensitivity, diabetes, gastro-oesophageal reflux disease, smoking status and primary-versus-revision surgery). These tests were also repeated for CRS only subgroup and showed no significantly differentially abundant genera across the different covariate levels.

**Figure 4:**
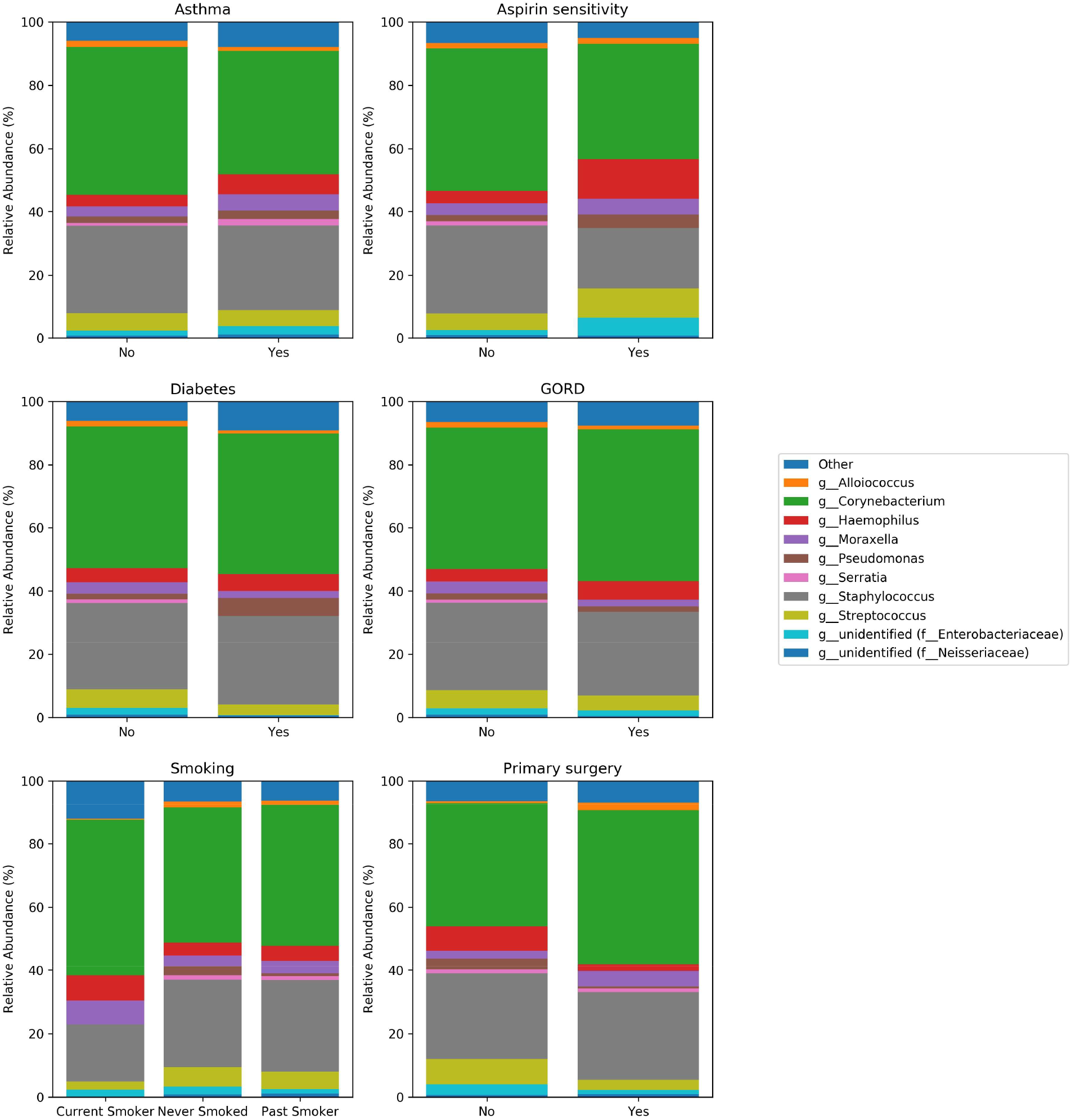
Relationship between clinical co-variates and microbial composition. Relative abundances of patient cohort (n = 410) when grouped by clinical co-variates (asthma status, aspirin sensitivity, diabetes, gastro-oesophageal reflux disease, smoking and surgery status). GORD = gastro-oesophageal reflux disease.

## DISCUSSION

The present study characterises the sinonasal microbiome in a large cohort of subjects from centres around the world using 16S rRNA surveillance. By adopting a unified, consistent methodology from sample acquisition to analysis we have been able to address many of the current existing limitations of currently available data.

We have identified *Corynebacterium, Staphylococcus, Moraxella, Streptococcus* and *Haemophilus* as the most abundant genera within the middle meatus of patients with or without CRS. This consistent finding across disease state and geography suggests that these organisms may form the core microbiome within the sinonasal tract. Interestingly, *Streptococcus, Moraxella* and *Haemophilus* species are traditionally respiratory tract organisms and constitute the most commonly cultured pathogens in patients with acute bacterial tonsillitis, otitis media and acute sinusitis.^13,33–35^ Anatomically, the sinonasal tract connects these three distinct anatomical regions and while these organisms appear to be commensals, so it is possible that the sinuses act as a reservoir for these organisms to subsequently initiate acute infection. While relatively low bacterial burden is present within the sinonasal mucosa, a dysbiosis within the diseased state may lead to over-representation of these organisms that subsequently lead to acute infectious or inflammatory processes.

While variations across geography have been demonstrated in the gut microbiome^1^ and have been suspected within the sinuses,^36^ this is the first study to examine this using standardized methodology. Our results suggest that although the bacterial composition of the core microbiome was preserved across the different sites, significant differences in both alpha and beta diversity occurred according to geography. Furthermore, microbiome composition also varied between centres. Most centres demonstrated a similar consistency in relative abundances of *Corynebacterium* and Staphylococcal species. Some centres, however, appeared to exhibit a varying microbial ecology of the less abundant organisms, which were either over- or under-represented. The most distinct microbial distribution was observed in samples collected from the Amsterdam centre, which appeared to be the most distinct centre with depletion in *Corynebacterium* and over-representation in Staphylococcus. Interestingly, antibiotic use in The Netherlands is amongst the lowest in the developed world.^37^ Such practices may influence the microbiota through selective microbial suppression and could partially account for the unique microbiome observed in Amsterdam. Overall, these findings suggest that there are differences in the sinonasal microbiome composition across geographical regions, but these differences would not alter the general pattern of core organisms described above. The driving factors behind geographical variations are yet to be elucidated - possible influences are diet, lifestyle, antibiotic prescribing patterns or environmental exposures.

Previous 16S rRNA gene sequence analyses of sinus microbiota have detected and ascribed importance to organisms such as *Cyanobacteria, Bacteroidetes, Propionibacterium* and *Acinetobacter*.^25,27–29,38^ These organisms were either not encountered or detected only in minute abundances in our analysis and given their unexpected nature in the nasal cavity, they could represent a source of contamination (either during sample collection or from DNA extraction reagents) or artefacts from bioinformatics pipelines. *Cyanobacterium*, for example is an environmental organism and *Bacteroidetes* is typically associated with the colon.^39–41^ In light of the limitations of current methodologies, reports of airway organisms that are novel, rare, and that do not form a part of the core microbiome, should therefore be interpreted cautiously. It is expected that the resolution of identification and functional characterization of bacterial species and strains in the sinuses can only improve, with advancement in sequencing technologies.

The organisms *Corynebacterium* and *Staphylococcus* are of particular interest. It was interesting to note that these organisms remained the two most abundant genera across geographical divides. Furthermore, amongst our control cohort, they were present in almost all individuals, again suggesting a key commensal function in the healthy state. Our findings of great abundance in both healthy patients and CRS patients mirrors previous studies.^22^ While *Corynebacterium* has traditionally been thought of as a nasal commensal,^42^ some studies suggest that certain species such as *C. tuberculostearicum* may be involved in CRS pathogenesis.^43^ We could not resolve the *Corynebacterium genus* in our current analysis to the species level, given the limitations of short-read 16S rRNA gene sequencing, but it is likely that a number of different corynebacterial species reside within the nose – the majority as commensals. Recent evidence shows that certain species of *Corynebacterium* are beneficial in the nasal airways. *Corynebacterium accolens*, for example, is a common nasal colonizer, and can inhibit streptococcal growth by releasing oleaic acids through hydrolysis of host triacylglycerols.^44^ In contrast staphylococcal species, and particularly *S. aureus*, by contrast have typically been viewed as potential pathogenic organisms within the nose. While being an asyptomatic coloniser in some individuals,^45^ *S. aureus* contributes to severe antibiotic- and surgery-resistant CRS.^46–48^ Our confirmation of high prevalence and relative abundance of *Staphylococcus* amongst controls suggests a role in maintaining a healthy sinus. Poor resolution of 16S analysis beyond the genus level results in both commensal and potentially pathogenic *Staphylococcus* species being grouped together. The outcome is that the healthy sinonasal microbiome may be composed of coagulase-negative commensal staphylococcal species, *S. aureus*, or more likely, a combination of both. If *S. aureus* is present in high abundance within the healthy sinus then perhaps this species plays a dual role within the sinuses: being essential for normal function typically, but sometimes becoming virulent at times of disease. The trigger for such a switch in roles is yet to be clarified but could be explained by the presence of *Corynebacterium*. Ramsey et al. described the ability of *C. striatum* to shift the gene profile of *S. aureus* away from virulence and towards commensalism.^49^ This hypothesis would be supported by our result demonstrating reduced *Corynebacterium* amongst CRSwNP. The depletion of these organisms may allow *S. aureus* to switch on virulence genes, propagating disease. Interestingly, CRSwNP is a more severe and resistant form of disease and has previously been linked with S. aureus virulence factors and, in particular, superantigens.^50,51^ Future studies and techniques which are able to identify microbiota composition to the species or strain level are required to further our understanding of these processes. Ultimately, the results from both current and previous findings highlight the complexity of the sinonasal microbiome and the myriad of functions (both human and bacterial) that act in concert to maintain homeostasis or produce disease. While we have presented data that has solidified our understanding of the upper airway microbial ecology, investigations into the microbiome function or metatranscriptome may provide further novel perspectives on important and critical microbial and host pathways.

## CONCLUSION

Understanding the characteristics of the sinonasal microbiome may provide novel insights to the normally functioning upper airway. This may increase our understanding of the pathobiology of diseases such as CRS. This study is the largest yet to describe the sinus microbiome and the first to examine geographical variation. We demonstrate that the core microbiome is composed of *Corynebacterium, Staphylococcus, Streptococcus, Moraxella* and *Haemophilus*. These organisms are present across disease phenotypes and countries. Utilising a large cohort and standardised methodology has allowed us to better characterise the sinonasal microbiome. By doing so, we have presented a foundation for future prospective studies into pathological states as well as functional analysis.

## METHODS

### Participating centres

A total of fourteen centres participated in the completion of this investigation. Thirteen centres provided patient samples for utilisation within the study. One centre (Northern Arizona University) provided bioinformatics expertise and consultation. The project was approved by the respective institutional human research ethics boards of all sample-collection centres (Table S1). Participating centres are listed below:

1. University of Adelaide, Adelaide, Australia (Lead Centre)
2. University of Sydney, Sydney, Australia
3. University of Auckland, Auckland, New Zealand
4. Siriraj Hospital, Mahidol University, Bangkok, Thailand
5. LV Prasad Institute, Hyderabad, India
6. University of Sao Paulo, Sao Paulo, Brazil
7. Catholic University of Chile, Santiago, Chile
8. Academic Medical Centre, Amsterdam, The Netherlands
9. McGill University, Montreal, Canada
10. Stanford University, California, USA
11. Harvard Medical School, Boston, USA
12. Medical University of South Carolina, Charleston, USA
13. University of Texas, Texas, USA
14. Northern Arizona University, Arizona, USA (Bioinformatics expertise)

### Study design

This study was a multi-centre, international collaborative investigation with a prospective, cross-sectional design. In total, thirteen centres across 9 countries provided samples for analysis. Written consent to tissue and clinical data collection was procured from all participants prior to surgery. Collection was performed during either endoscopic sinonasal surgery or ancillary otolaryngological procedures, such as tonsillectomy, septoplasty or skull base tumour resections. Individuals were classified as having CRS if they fulfilled the criteria outlined in the International Consensus statement on Allergy and Rhinology: Rhinosinusitis.^11^ Further sub-classification according to the absence (CRSsNP) or presence (CRSwNP) of nasal polyps was performed by endoscopic assessment at the time of surgery. Patients who underwent ancillary otolaryngological procedures and had no clinical or radiological evidence of CRS formed the healthy control cohort.

### Metadata collection

Clinical data was collected using standardized patient questionnaires undertaken by each participant at the time of study consent. In non-English speaking countries, the questionnaire was translated by a qualified interpreter and checked for accuracy by the lead investigator from that country. Collected data included patient demographics, medical history, ethnicity, social and environmental exposures as well as quality of life scoring via the validated SNOT-22 and Visual Analogue Scores (VAS). The standardized questionnaires used (which we termed the Open Source Sinonasal Survey “OS3”) are freely available at https://github.com/adelaide-orl/os3.

### Sample collection and transport

Each centre was asked to provide microbiome swabs from non-CRS controls and CRS patients for analysis. Patients were all anaesthetised at the time of sample acquisition. All samples were collected in a standardized manner prior to commencement of any surgical intervention, whilst the nasal cavity remained unaltered. Microbiome swabs were collected intra-operatively using guarded and endoscopically-guided Copan Flocked swabs (COPAN ITALIA, Brescia, Italy) to sample the middle meatus.^24,52^ Swab heads were subsequently separated into sterile cryotubes (Sarstedt, Numbrecht, Germany), placed on ice immediately, and then transported to −80°C storage. Once all samples had been acquired from a centre, they were then transported to the lead centre (Adelaide) using a secure cold chain (Cryoport, Irvine, CA, USA). This was to ensure standardized down-stream processing. The standard Cryoport containers used for shipment of samples are liquid nitrogen dewars capable of keeping stored samples at a stable temperature for at least 15 days. These containers have continuous temperature monitoring to ensure preservation of the cold chain throughout the shipment. Any evidence of temperature disturbance, or displacement or damage to transported cryotubes resulted in those samples being excluded from further processing.

### DNA Extraction

All DNA extraction was performed at the lead centre using Qiagen DNeasy Blood and Tissue Kit (Qiagen, Hilden, Germany), as per manufacturer’s instructions. Total DNA was extracted from all clinical samples as well as a DNA-negative control with extraction reagents only. All extractions were undertaken in strict sterile conditions, utilising new equipment for each sample to exclude cross-contamination. In brief, swab heads were prepared for extraction by being cut to 2-3mm pieces and placed into a microcentrifuge tube. A lysozyme (Sigma, St. Louis, USA) solution at 20mg/ml in Lysis buffer (20 mM Tris-Cl, pH8; 2mM sodium EDTA; 1.2% Triton X-100, filter sterilised; Sigma, St Louis, USA) was added to each sample and left overnight at room temperature. Samples were then homogenised using 5mm steel beads and a Tissue Lyser II (Qiagen, Hilden, Germany) at 15Hz for 20 seconds. Steel beads were then removed prior to further homogenisation with 50mg glass beads, again using the Tissue Lyser II at 30Hz for 5 minutes. Proteinase K and Buffer AL (Qiagen, Hilden, Germany) were added to each sample and left to incubate for 30 minutes at 56°C. Tubes were then centrifuged briefly to collect beads and supernatant transferred to a fresh microcentrifuge tube. After addition of 100% ethanol to supernatant, the new mixture was pipetted into DNeasy Mini Spin columns (Qiagen, Hilden, Germany). Subsequent extraction of DNA from supernatant mixture was as per manufacturer instructions, with a total of 100ul of DNA extracted per sample. Concentration was determined using a NanoDrop 1000 Spectrophotometer (Thermo Scientific, Massachusetts, USA). Extracted DNA was stored at −80°C until sequencing. Any samples that were suspected to be improperly handled, contaminated or did not pass quality control during processing were excluded. A total of 532 samples (126 CRSsNP; 212 CRSwNP; 194 Controls) passed all stages of transport and processing to be sent to the sequencing facility (Australian Genome Research Facility; AGRF, Melbourne, Vic, Australia).

### Polymerase Chain Reaction (PCR) Amplification of the 16S rRNA gene and sequencing

PCR and sequencing were performed by AGRF. Libraries were generated by amplifying (341F–806R) primers against the V3–V4 hypervariable region of the 16S rRNA gene. (CCTAYGGGRBGCASCAG forward primer; GGACTACNNGGGTATCTAAT reverse primer).^53^ PCR was done using AmpliTaq Gold 360 master mix (Life Technologies, Mulgrave, Australia) following a two-stage PCR protocol (29 cycles for the first stage; and 14 cycles for the second, indexing stage). Concentrations of the resulting amplified amplicons were measured using fluorometry (Invitrogen Picogreen; Thermo Fisher Scientific, Waltham, MA, USA). Amplicons were normalised according to the obtained concentrations prior to sequencing. Sequencing was done on the Illumina MiSeq platform (Illumina Inc., San Diego, CA, USA) with the 300-base-pairs paired-end chemistry over 8 runs.

### Bioinformatics pipeline

Demultiplexed fastq files were received from the sequencing facility. We used the new QIIME 2 (version 2018.11)^30^ for our bioinformatics pipeline, utilizing various QIIME 2 plugins at each step. Forward and reverse reads were joined using PEAR^54^ through the QIIME 2 plugin q2-pear (https://github.com/bassio/q2-pear). Joined sequences were then quality-filtered using the QIIME 2 plugin q2-quality-filter,^55^ with minimum quality 20, according to recommendations.^56^ This was followed by abundance-filtering applied on the reads, according to the method by Wang et al.^57^, through the python implementation in the QIIME 2 plugin q2-abundance-filtering (https://github.com/bassio/q2-abundance-filtering). Denoising and Amplicon Sequence Variant (ASV) formation were done using deblur^56^ through the q2-deblur plugin using the parameters (trim-size = 435; min-size = 1; min-size = 1). Taxonomy assignment was done against the Greengenes 16S reference database (the 99% clustered similarity sequences),^58^ version 13.8 (August 2013) using the BLAST-based classifier implemented in QIIME 2 (q2-feature-classifier)^59^ and which implements a Lowest Common Ancestor (LCA) consensus algorithm. To address limitations of de novo trees generated from short-length ASVs, we utilized the SATé-enabled phylogenetic placement (SEPP) technique^60^ for insertion of the ASVs into the high-quality tree generated from the 99% OTUs Greengenes reference database, and the ASVs that did not fit anywhere into the tree were filtered out of the ASV table.

A rarefaction depth cut-off was chosen at 400 before downstream diversity analysis and comparisons of relative abundances of taxa. Alpha rarefaction plots of unique number of ASVs in each sample (i.e. richness), as well as Shannon’s diversity index confirmed almost all samples reaching a plateau at this depth indicating sufficient sampling. (Supplementary Figure S2) Applying this depth yielded 410 (out of 532) samples for downstream analysis. Taxa were mostly compared at the genus level. Mean relative abundance as well as prevalence of the genera were calculated for each group. Faith’s phylogenetic diversity index^31^ was used for alpha diversity and weighted Unifrac^32^ distance matrices were calculated for beta diversity analyses. Diversity metrics were generated through sci-kit bio version 0.5.3.

### Statistical analysis

Statistics were done using packages from the Python Scientific Stack^61^ and R (R Foundation for Statistical Computing, Vienna, Austria) through the Jupyter notebook interface^62^, utilizing the assistance of packages from the Scientific Python^61^ stack (numpy, scipy, pandas, statsmodels), scikit-bio (https://github.com/biocore/scikit-bio) and omicexperiment (https://www.github.com/bassio/omicexperiment).

We investigated the relative abundances of genera in different subgroups using linear mixed model analysis (R packages “lme4” and “lmerTest”). Linear mixed effects modelling was performed to control for the “centre” variable, which was included in the model as a random effect. The mixed models were fit using the restricted maximum likelihood (REML) as implemented in the default method in the “lme4” package. Mean fixed effects of variables were extracted from the model objects in R using the R package “emmeans”, (the successor to “lsmeans”).^63^ The p values for the covariates in the mixed models were generated using t-tests using Satterthwaite’s method as implemented in the “lmerTest” package.^64^

Comparison of mean relative abundances of the top 10 taxa between centres, and comparison of mean alpha diversity indices between disease groups and centres, were performed using Mann-Whitney-Wilcoxon tests, with multiple comparisons correction using the Benjamini-Hochberg method.^65^ For multivariate analysis of beta diversity metrics, we employed permutational multiple analysis of variance (PERMANOVA)^66^ implemented in the function “adonis” from the R package “vegan”.^67^

## Funding information, Disclosures and Conflicts of Interest (COI)

Mohammad Javed Ali:

Receives royalties from Springer for his treatise “Principles and Practice of Lacrimal Surgery” and “Atlas of Lacrimal Drainage Disorders”.

No conflict of interest relevant to this study.

Ahmed Bassiouni, Clare Cooksley, Mahnaz Ramezanpour, Sophia Moraitis:

No conflict of interest to declare.

Benjamin Bleier:

Grant Funding: R01 NS108968-01 NIH/NINDS (Bleier PI) – This isn’t relevant to this study.

Consultant for: Gyrus ACMI Olympus, Canon, Karl Storz, Medtronic, and Sinopsys.

Equity: Cerebent, Inc, Arrinex.

COI: None relevant to this study.

Claudio Callejas:

No conflict of interest to declare.

J Gregory Caporaso, Matthew R Dillon, Arron Shiffer:

No conflicts of interest to declare. This work was funded in part by National Science Foundation Award 1565100 to JGC.

Emily K Cope:

Financial information: This work was partially funded under the State of Arizona Technology and Research Initiative Fund (TRIF), administered by the Arizona Board of Regents, through Northern Arizona University.

No relevant disclosures or COI.

Marjolein E Cornet:

No financial relationships or sponsors. No conflicts of interests.

Richard G Douglas:

Received consultancy fees from Lyra Therapeutics and is a consultant for Medtronic. These are not relevant to this study.

Daniel Dutra:

No conflict of interest to declare.

Christos Georgalas:

No conflicts of interest to declare.

Richard J Harvey:

Consultant with Medtronic, Olympus and NeilMed pharmaceuticals. He has also been on the speakers’ bureau for Glaxo-Smith-Kline, Seqiris and Astra-Zeneca.

No direct conflict of interest to declare.

Peter H Hwang:

Financial Relationships: Consultancies with Arrinex, Bioinspire, Canon, Lyra Therapeutics, Medtronic, Tivic.

Conflicts of Interest: None.

Amber U Luong:

Serves as a consultant for Aerin Medical (Sunnyvale, CA), Arrinex (Redwood City, CA), Lyra Therapeutics (Watertown, MA), and Stryker (Kalamazoo, MI) and is on the advisory board for ENTvantage (Austin, TX).

Her department receives funding from Genetech/Roche (San Francisco, CA) and AstraZeneca (Cambridge, England).

No COI to declare related to this study.

Sathish Paramasivan:

Supported by a Garnett Passe and Rodney Williams Memorial Foundation Academic Surgeon Scientist Research Scholarship.

No conflicts of interest to declare.

Alkis J Psaltis:

Consultant for Aerin Devices and ENT technologies and is on the speakers’ bureau for Smith and Nephew. Received consultancy fees from Lyra Therapeutics. These are not relevant to this study.

Rodney J Schlosser:

Grant support from OptiNose, Entellus, and IntersectENT (not relevant to this study). Consultant for Olympus, Meda, and Arrinex (not relevant to this study).

Pongsakorn Tantilipikorn:

No financial disclosures or conflict of interest.

Marc A Tewfik:

Principal Investigator: Sanofi, Roche/Genentech, AstraZeneca.

Speaker/Consultant: Stryker, Ondine Biomedical, Novartis, MEDA, Mylan.

Royalties for book sales: Thieme.

Sarah Vreugde:

No conflicts of interest relevant to this study.

Peter-John Wormald:

Receives royalties from Medtronic, Integra, and Scopis, and is a consultant for NeilMed. These are not relevant to this study.

## Abbreviations

NGS: Next-generation sequencing
CRS: chronic rhinosinusitis
CRSsNP: chronic rhinosinusitis sans nasal polyps
CRSwNP: chronic rhinosinusitis with nasal polyps

## Supplementary Tables

**Table S1:** Ethics approvals.

| Centre | Approval |
| --- | --- |
| University of Adelaide,<br>Adelaide, Australia | The Queen Elizabeth Hospital Human Research Ethics Committee<br>(approval HREC/14/TQEHLMH/222) |
| University of Sydney, Sydney,<br>Australia | HREC Reference number: LNR/14/SVH/373 |
| University of Auckland,<br>Auckland, New Zealand | HDC Northern Regional Ethics Committee approval<br>NTX/08/12/126/AM13 |
| Siriraj Hospital, Mahidol<br>University, Bangkok , Thailand | Siriraj Institutional Review Board: Certificate of Approval 246/2559<br>(EC1) |
| LV Prasad Institute, Hyderabad,<br>India | Local ethics approval number LEC-01-15-001 |
| University of Sao Paulo, Sao<br>Paulo, Brazil | Local ethics approval number CAAE: 46421415.8.0000.0068 |
| Catholic University of Chile,<br>Santiago, Chile | Scientific Ethics Committee, Faculty of Medicine, Pontifical Catholic<br>University of Chile (reference number 14-435) |
| Academic Medical Center,<br>Amsterdam, The Netherlands | METC number: W14_282 # 14.17.0338 |
| McGill University, Montreal,<br>Canada | McGill University Health Centre, Biomedical D Research Ethics<br>Board, Study Number 14-433 BMD |
| Stanford University, California,<br>USA | Ethics Protocol ID : 33326 |
| Harvard Medical School, Boston,<br>USA | Massachusetts Eye and Ear IRB Approval #651070-9 |
| Medical University of South<br>Carolina, Charleston, USA | Ethics/Institutional Review Board: Pro00038979 |
| University of Texas, Texas, USA | Ethics approval HSC-MS-16-0853 |

**Table S2A:**
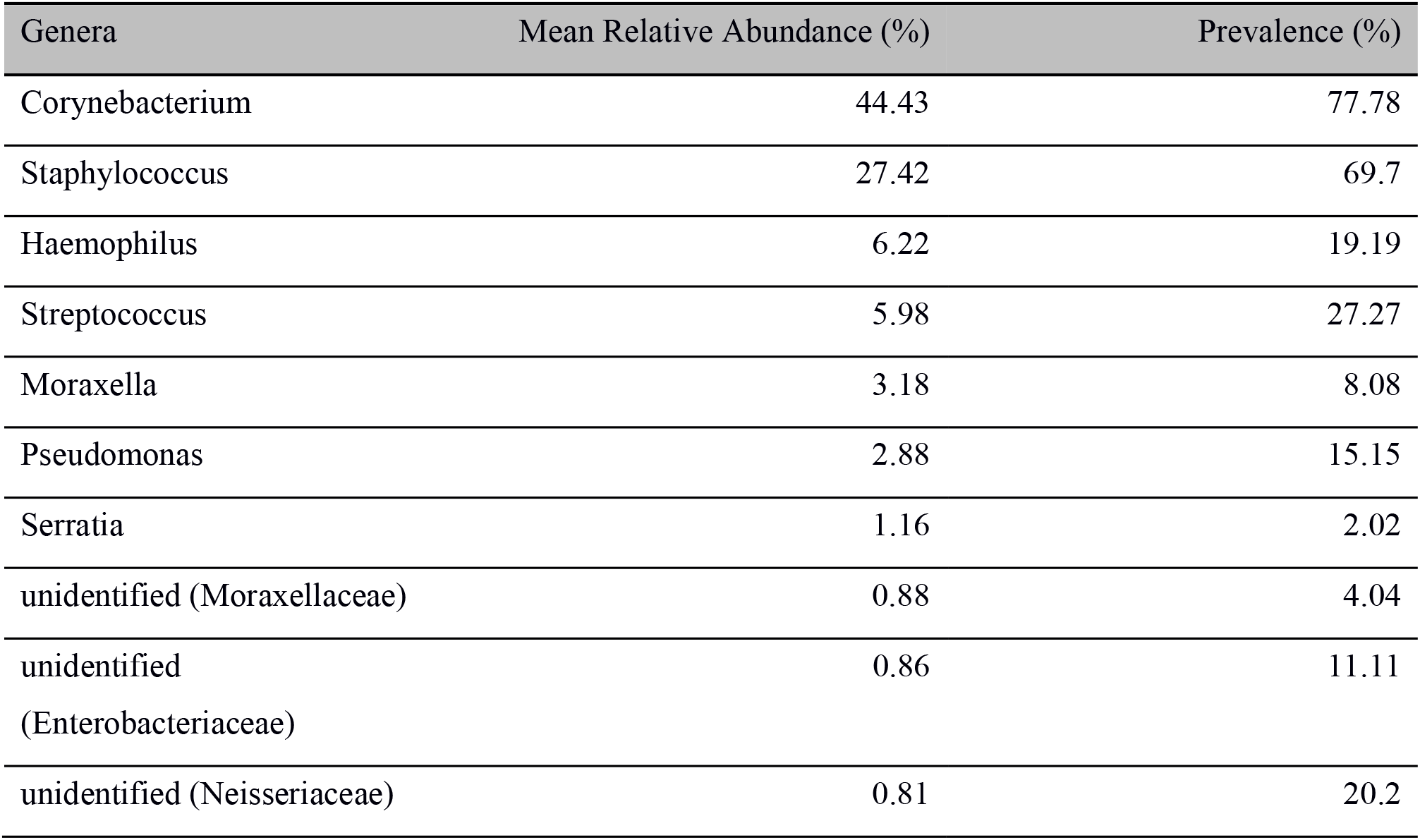
Abundance and prevalence of genera in CRSsNP patients.

| Genera | Mean Relative Abundance (%) | Prevalence (%) |
| --- | --- | --- |
| Corynebacterium | 44.43 | 77.78 |
| Staphylococcus | 27.42 | 69.7 |
| Haemophilus | 6.22 | 19.19 |
| Streptococcus | 5.98 | 27.27 |
| Moraxella | 3.18 | 8.08 |
| Pseudomonas | 2.88 | 15.15 |
| Serratia | 1.16 | 2.02 |
| unidentified (Moraxellaceae) | 0.88 | 4.04 |
| unidentified<br>(Enterobacteriaceae) | 0.86 | 11.11 |
| unidentified (Neisseriaceae) | 0.81 | 20.2 |

**Table S2B:**
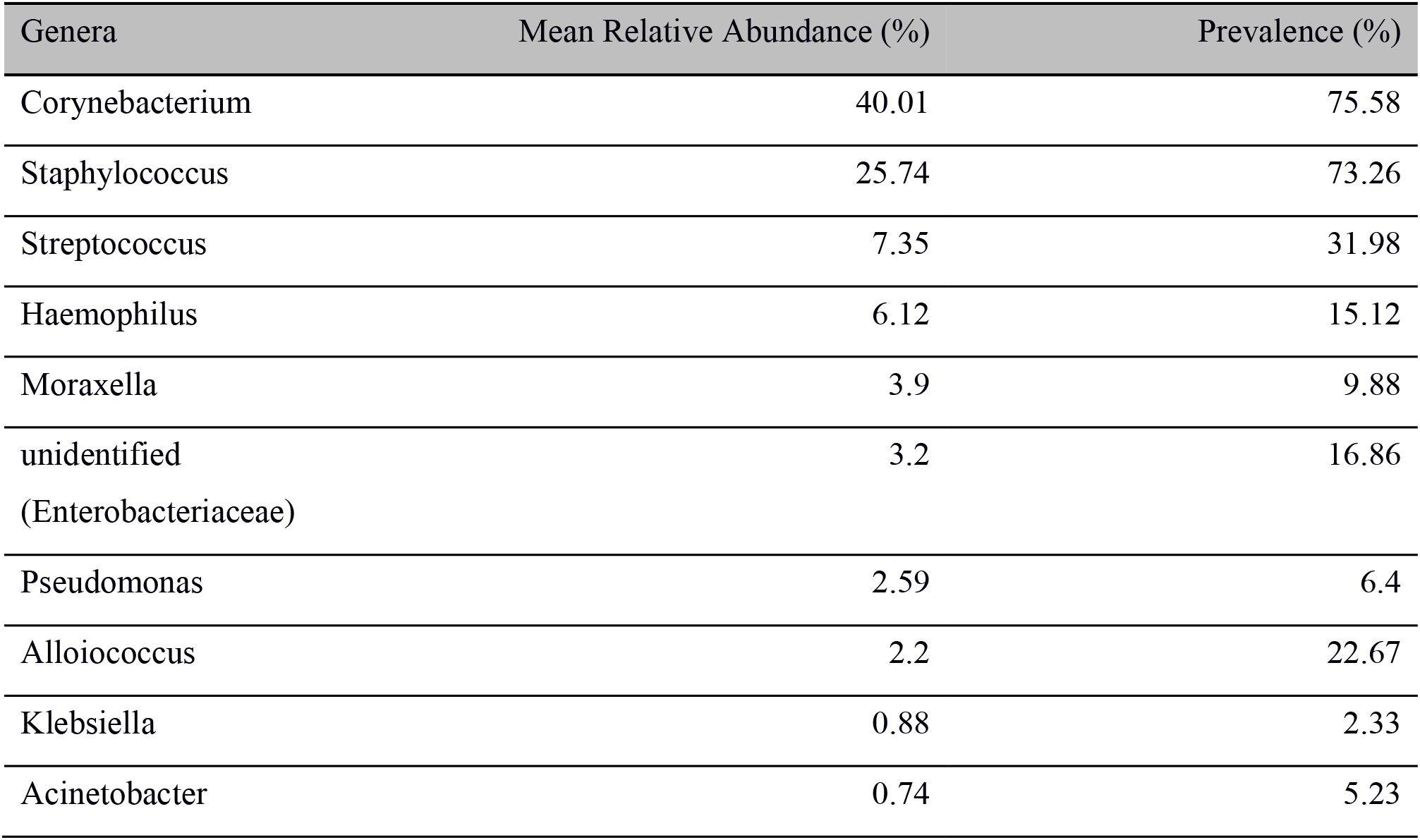
Abundance and prevalence of genera in CRSwNP patients.

## Supplementary Figures

**Figure S1:**
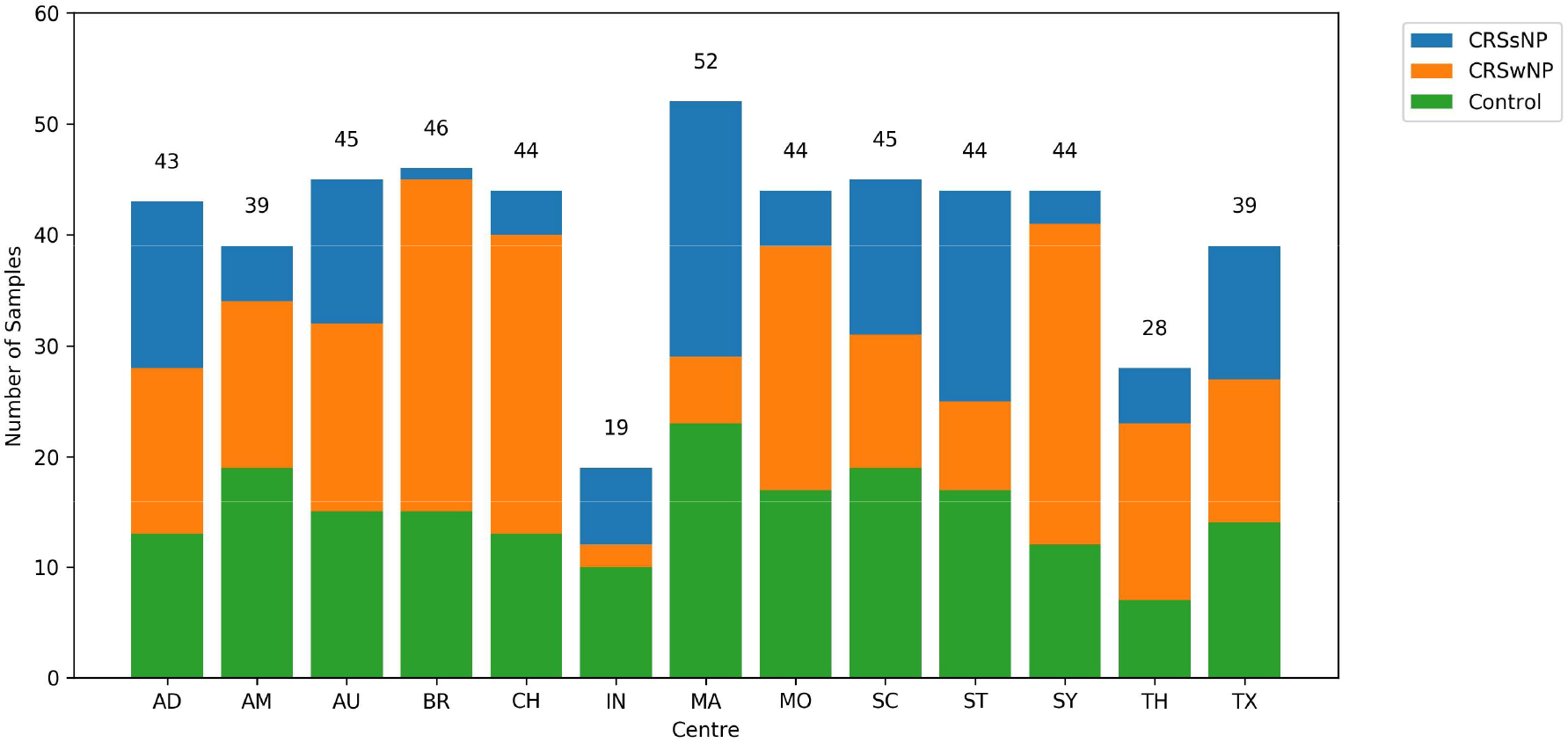
Collected sample disease characteristics by centre. Figure demonstrating the disease characteristics of patients from each centre. Each bar represents cumulative numbers. Control patients are indicated by, CRSsNP by blue and CRSwNP by orange. Final contributing numbers are indicated at the peak of each bar. AD = Adelaide; AM = Amsterdam; AU = Auckland; BR = Brazil; CH = Chile; IN = India; MA = Massachusetts; MO = Montreal; SC = South Carolina; ST = Stanford; SY = Sydney; TH = Thailand; TX = Texas.

**Figure S2:**
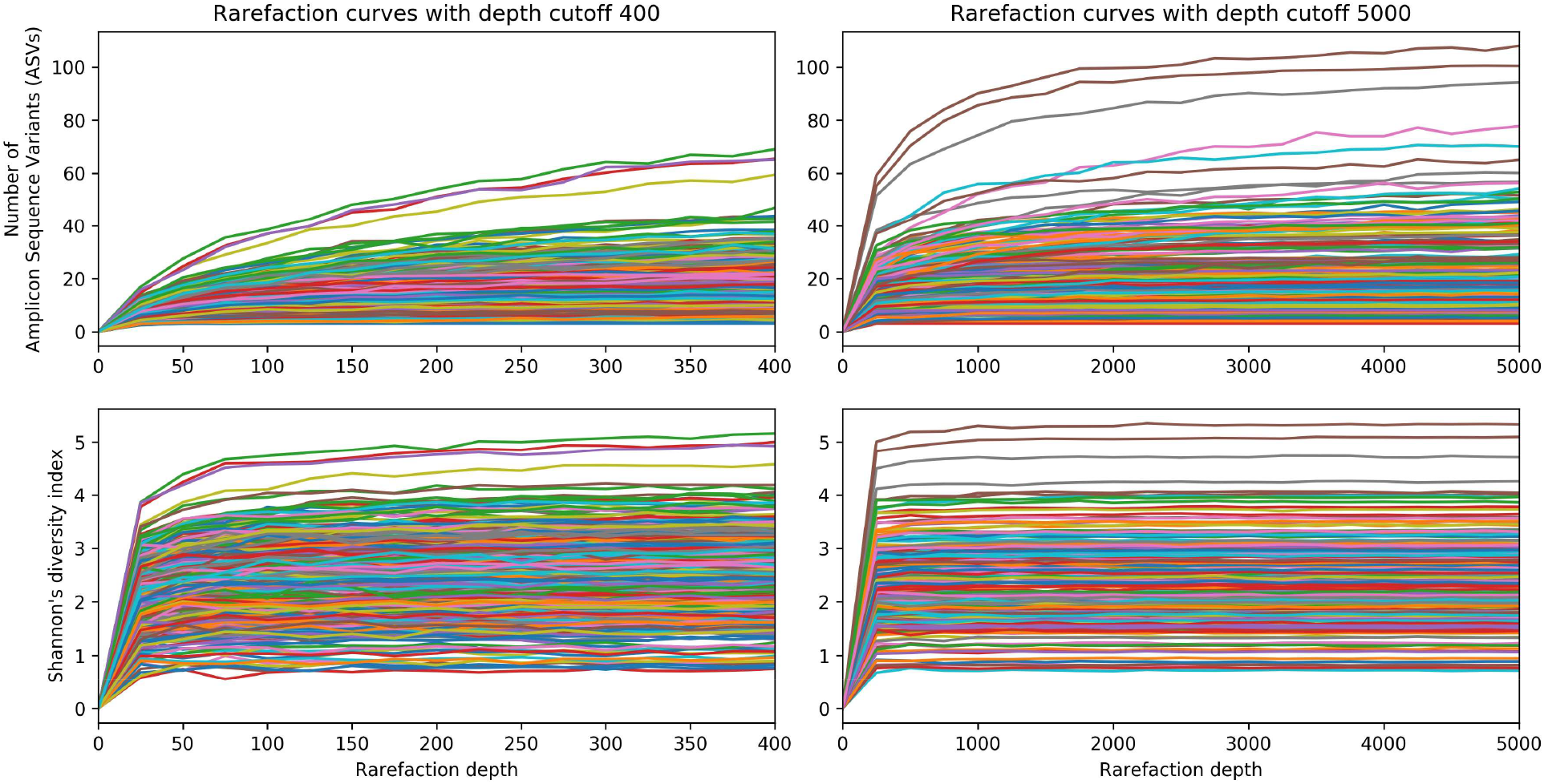
Rarefaction analysis: Rarefaction curves plot. A rarefaction depth of 400 was chosen before downstream diversity analysis and comparisons of relative abundances of taxa. Rarefaction plots (at depth cutoffs 400 and 5000) of richness (total number of ASVs per sample) and Shannon’s alpha diversity index confirmed almost all samples reaching a plateau at this depth indicating sufficient sampling. Each line in the plot represents a single sample. ASVs = Amplicon Sequence Variants.

**Figure S3:**
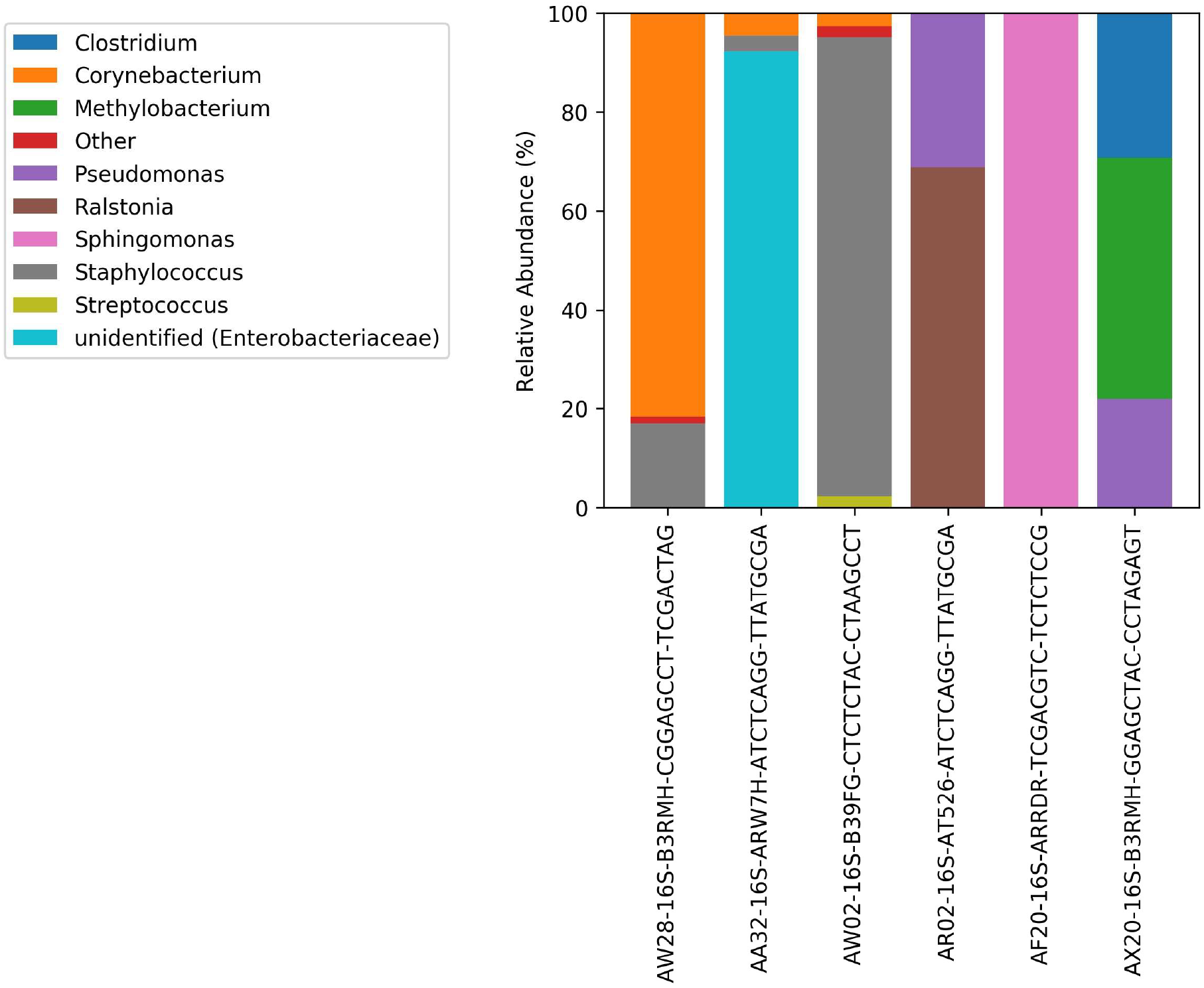
Taxa bar plots of the negative template control samples (i.e. DNA-negative samples with extraction reagents only). The rightmost three (of the six) samples were below the rarefaction threshold depth of 400 reads.

